# HUB-DT: A tool for unsupervised behavioural discovery and analysis

**DOI:** 10.1101/2024.01.08.574696

**Authors:** Adrian Lindsay, Jeremy K Seamans

**Affiliations:** Djavad Mowafaghian Centre for Brain Health, Department of Psychiatry, University of British Columbia

## Abstract

There has been an expansion in the diversity of tools used to measure various aspects of brain function in behaving animals. While these tools have great potential to transform our understanding of brain function, they are of little value if the behavior of interest is poorly defined or quantified. Traditional methods of behavioural labelling focus on easily quantified gross measure, such as velocity, gate crossing, nosepokes, etc. While these measures are specific and reproducible, they are crude descriptions of behaviour at best. Manually defined behaviours, while providing increased granularity and descriptive power over specific gross measures, suffer from being inexact and somewhat arbitrary. Consistent labelling between human observers is often difficult, and even if manually defined behaviours are subsequently labelled in an automated fashion (via a supervised learning algorithm) these behaviours need to be defined ahead of time, possibly biasing the range of behaviours of interest for a given task.

Here we present HUB-DT, a behavioural discovery pipeline built on the frameworks of several existing tools and methods in the space of behavioural categorisation, the specifics of which will be highlighted in this report, and designed to address the requirements of behavioral discovery.

## Introduction

While unsupervised behavioural categorisation is relatively recent to the field of neuroscience, there is a rapidly growing body of tools and techniques to use. DeepLabCut (Mathis et al, 2018) is now commonly used to define the positions of body parts of interest in each frame of a video, providing easy limb tracking through time. While a static set of limb coordinates is not behavior, various tools have been developed to translate limb positions through time into behavioral categories. DeepBehavior: Automated Analysis of Behavioural Kinematics (Arac et al 2019). VAME: Segmenting behaviour using latent space representations and a HMM (Luxem et al 2020). MoSeq: Mapping sub-second structure in mouse behaviour (Wiltschko et al 2015). B-soid: Unsupervised behaviour identification (Hsu et al 2021), and Motion Mapper: Mapping stereotyped behavior in fruit flies (Berman et al 2014), to name just a few. Each approach has advantages and disadvantages, these will be explored in more detail below, that the user must consider with regards to their specific application and question.

‘Behaviour’ in this context is often treated in a general and nonspecific manner, and what defines a behaviour can depend on the particulars of the approach. We would advocate for an approach that leverages the power of machine learning to exhaustively characterize all the fine motor movements produced under various, closely-related conditions. One drawback is that this is only possible by tracking the fine movements of multiple limbs, which necessitates either incorporating inputs from multiple camera views to avoid occlusions of a limb in any given frame, utilizing 3D depth imaging or motion capture, or using a model to infer the occluded features.

An additional concern is how to balance the desire for ‘a priori free’ behavioral detection with interpretability of the detected behaviors. To address this issue, one should be able to directly compare automated behavioral labels with human curated behavioural labels in a common space or frame of reference. But, even if automated and human curated behavioral labels agree, the behavior may not be what it appears. Consider for example the case of fear conditioning, a commonly used task used to assess fear memory in rodents. Freezing in response to a conditioned cue is taken to be evidence of fear learning, but how can one be sure the animal is freezing because they recalled a fearful memory versus just simply not moving? One must therefore consider whether the behavior is actually biologically relevant to the animal. One route to assess whether the animal may view a detected behavior as discrete or meaningful would be to check whether a specific pattern of neural activity emerges whenever the labelled behaviour is detected. To facilitate this type of comparative analysis, one should also be able to directly compare behavioral labels with neural data in a common frame of reference.

Even in cases where an existing tool or method is a good fit for analysis on a task, there is always tuning and parameterization required. This can be significant in terms of time investment. Indeed, in practical terms, the importance of usability, modifiability, and computational cost and efficiency in the design and adoption of a particular tool or technique cannot be overstated. It is important to tailor the analysis methods to the needs of the user and the goals of the study.

As mentioned, our novel behavioural discovery pipeline, HUB-DT, was built to provide unsupervised behavioural classification in light of the constraints discussed above. It was also was designed to address our specific use cases that typically involve rats performing free-ranging behaviours in relatively unconstrained environments while we record the activity of tens to hundreds of neurons in the frontal cortex. One challenge that is somewhat unique to the frontal cortex is that these neurons are extraordinarily multi-modal and flexible in terms of how they encode seemingly static events in the world (see Seamans and Floresco 2022 for review). As a result, HUB-DT was designed to provide extreme flexibility in the way behaviours are discovered while also incorporating some basic analysis techniques that we have found to be effective in connecting free-ranging behaviors to the neural ensemble activity.

HUB-DT can perform traditional behavioural annotation as well as ‘parameter-free’ behavioural discovery in discreet steps, maintaining the connection to human-sensible categories and interpretability. It applies the same basic approach utilized by Motion Mapper, fitting wavelets to the frequency components of individual limb movements. It then employs various clustering techniques (shared across several of the discussed tools) to find the unique constellations of limb movement frequencies that consistently arose within a session (i.e. “behaviours”). This tool enables us to discover and label behaviours in our experimental tasks, and to perform comparative analysis with manually annotated behaviours as well as simultaneously recorded electrophysiology data.

What follows is a description of the construction and function of the HUB-DT tool, as well as a brief exploration of the motivation behind its’ features and development. We will briefly describe how to manually track annotated behaviours and how it can be used for unsupervised behavioral discovery. HUB-DT allows the user to discovery behaviours at varying levels of granularity and to compare discovered behaviours to manually labelled behaviours in a common space. We will highlight how the unsupervised behavioural discovery process is able to detect subtle differences in behaviours that are unfeasible for human observers to reliably annotate, and how the model and parameters can be adjusted to provide the best correspondence between neural activity and discovered behaviours. We will also discuss the in-built methods for interrogating the behaviours detected by HUB-DT, as well as methods for comparative analysis with neural data. We will also provide a brief overview of the tools’ graphical user interface, its’ features as well as a summary of its’ use and practical considerations. A more complete user guide will be available accompanying the HUB-DT Git repository.

To provide practical illustrations of how HUB-DT works, we will describe how it was used to identify the behaviours of rats engaged in a conditioned-outcome task where outcomes were either appetitive, aversive or neutral in different blocks of trials (Caracheo et al 2018; Lindsay et al 2018; 2023). It was able to detect block-specific behaviors and to divide neuronal activity into the portions related to discovered behaviors and the portions putatively related to the affective states evoked in each block of trials. We will also describe how HUB-DT was used to discover unique sets of behaviours expressed by rats as they transitioned through a morphine administration and withdrawal procedure. The results showed that some of the drug-associated behaviours detected by HUB-DT were those commonly used by human observers to define the various stages of addiction as well as other behaviours not previously described in the literature.

### A rich ecosystem of existing tools and techniques

There are a plethora of similar tools designed and built to extract and label behaviour, which at a surface level, pursue the same goal: automated behavioural annotation for use in experimental analysis. However, these tools are all implemented differently, have significant differences in function and scope, and as a result are optimal for the specific use cases and research goals for which they were designed. Here we include a brief survey of some of the existing methods, the basics of how they approach and achieve behavioural classification, and some use cases and observations.

#### DEEPBEHAVIOUR

(Arac et al 2019) is an end-to-end kinematic analysis pipeline, and represents an early approach in the field to using applied machine learning to extract behavioural data. Input comes from a multi-camera setup with high framerate and precise alignment, and a trained neural network (specifically a Convolutional neural network or long-short-term memory recurrent neural network) is used to provide markerless tracking of features from the input video. Features in this context are bounding boxes for limbs or digits of interest. These tracked features are then transformed into trajectories, which are clustered using a technique called dynamic time alignment kernels. This allows for flexible clustering, both supervised and unsupervised. DEEPBEHAVIOUR represents a combined computer vision approach, using different neural network architectures for analysis of different tasks and species. The framework enables in-depth trajectory analysis of these specific behaviours, and these identified kinematics provide the basis for downstream clustering/classification. This is a highly task specific approach. While the components can be more generally applicable, their use in an analysis pipeline requires considerable modification and tuning for any given task. In particular, training the Neural Networks and determining an appropriate space and algorithm for clustering/classification can be time and resource consuming.

#### Motion Mapper

(Berman et al 2014) is a spectrographic projection-based technique for identifying and analysing behavioural stereotypies, initially in fruit flies, and subsequently applied to some other lab animal species. As originally conceived, input came in the form of multiple camera views, which were aligned and used to perform postural extraction from raw video. These extracted postures are passed through a series of projection steps, where they are transformed into points in a high-dimensional behavioural space. This projection involves transforming the data into frequency space, and then multiplying the data by a series of morlet wavelets, resulting in spatial data quantized across a set of temporal ranges. This behavioural space is then reduced in dimension to 2-D (the remarkable degree to which information is preserved in as few as two dimensions is covered at length in the motion mapper technical paper) via a dimensionality reduction technique known as t-SNE (explained in greater detail below). This reduced behavioural space is then clustered via the watershed transform: the 2-d space is treated as a 3-d surface where height is determined by the density of points. This surface is iteratively filled at increasing depth, and the distinct watersheds that result from this filling are labelled as clusters.

Spectrographic projection is a key feature of this approach. This enables multi-resolution temporal dynamics to be represented in a combined space, encoding changes in pose segments (roughly equivalent to features as used in other tools) at multiple time scales. The resulting behavioural space is slightly biased towards periodic changes in pose within those frequency ranges selected as wavelets. In the pursuit of behavioural stereotypies, this can be beneficial, as one would expect stereotyped engrams to be at least somewhat periodic. This process of explicit postural extraction combined with spectrographic projection allows for in-depth trajectory analysis of behaviour space, i.e. location within the space through time gives a trajectory of dwelling and transitions between behaviours. Finally, the watershed transform as a means of clustering is essentially non-parametric, allowing for easy and consistent application. However, this can be an issue when the density space is less clearly separated: Specific to our application, fly behaviour is considerably more stereotyped and frequency-locked than rodent behaviour which results in a density space that is often poorly clustered via the watershed transform.

#### VAME

(Luxem et al 2020) uses variational autoencoders, a specific neural network architecture, to produce a behavioural latent space. These autoencoders are built on recurrent neural networks, which use fixed-length sequences of positional data (from annotated video, marker based tracking, or other means) as input. Autoencoders generate a latent space via a process of constrained representation: the network has limited neurons with which to represent the input space, which trains the network to compress and align the representation to accommodate this limited space. The behavioural space encodings that result from this form a lower dimensional manifold, representing behaviour. VAME then uses a set of Hidden Markov Models (HMMs) to generate state sequences within this behavioural space, which quantizes the dynamical system as a discrete-state continuous-time Markov chain.

This process is effective in identifying short behavioural motifs with common transitions such as phases of locomotion, progression in grooming, etc. The approach is appropriate for open-field use where subtle variations in these motif frequencies are of interest: investigating changes in walking patterns for example. VAME is however, relatively computationally expensive. Getting the technique up and running requires training of a set of neural networks and HMMs on any given data set, both of which require significant tuning. There are considerations resulting from the fixed length sequence input: the temporal signal is biased towards short fixed length sequences at both the autoencoder and HMM steps. The latent space is generated from a sizable complex neural network, and the state-sequence outputs are purely probabilistic. This combined complexity can make it difficult to interpret the contents and characteristics of a given output state.

#### B-SOID

(Hsu et al 2021) is a fixed-time unsupervised behavioural categorization tool designed to be used with rodents in open field experiments. The input space is generated from frame-difference measures applied to annotated video: the derivatives of position, angle, and relative pose (arrangement of tracked features) distance. This input set is generally fixed, and was selected by extensive information measure testing for the intended use case. This set of feature derivatives is then embedded in a reduced dimension space via the UMAP algorithm (covered in greater detail below), reducing it to two dimensions to allow the application of density-based clustering. A hierarchical density-based clustering algorithm called HDBSCAN (also explained in greater detail below) is then used to perform unsupervised clustering on this density space, resulting in a set of clusters as putative behaviours. The tool is, by default, setup for use with a single bottom-up view camera, but this is modifiable.

The critical component of this technique is the use of frame difference measures (the derivatives of position, angle, and relative pose) as the full input space. This enables the direct readout of kinematics from the output clusters, but means the clusters are defined across only a single fixed timescale. This timescale is dependent on framerate and the parameters of the frameshift performed by the tool, and performance can be impacted by noise and by slow framerates (by default, this is set for 50 fps). For reference, our use cases involve video at a range of framerates from 15 to 60 fps. The choice of input space and framerate tends to bias B-SOID towards short-sequence based behaviours: this is useful for data sets with large numbers of repeated sequences, as is common in many open-field applications, but less well suited for small datasets with high diversity where local structure may be more important. This can impact downstream behavioural selection, and may necessitate extensive modification if the input data is less suited to the design parameters.

An additional feature of the tool is propagating labelled behaviour across datasets. The creators of B-SOID demonstrated that training a Random Forest to reproduce clustering labels works well (this is also true of our approach, and we have adopted a similar procedure for propagating our discovered behaviours across datasets).

#### MoSeq

(Wiltschko et al 2015) is an unsupervised method designed to parse mouse behaviour into a set of behavioural syllables (usually sub-second behavioural motifs). Unique among the approaches mentioned here, MoSeq takes as input 3-D depth images/video, providing direct 3-D tracking without the need for multiple cameras. Subjects are extracted from the background, resulting in location agnostic 3-D models of single or even multiple mice. Similar to Motion Mapper, and indeed to our own tool, MoSeq utilizes wavelet projection to quantize behaviour across a set of temporal scales. MoSeq then uses a probabilistic time-series model, which they label an autoregressive hidden Markov model, or AR-HMM to segment behaviour into a set of behavioural motifs. In use, MoSeq provides a means to extract and label behavioural syllables, as well as extensive tools for analysing the transitions between behaviours which was the focus of the research which motivated the creation of the tool. Like all the techniques covered in this brief summary, there are some caveats and considerations to take into account with its use. Working with 3-D depth directly removes the need for calibration of multiple cameras, but does impose some limitations on the experimental space and data collection, particularly if electrophysiology or other data is being simultaneously collected. The AR-HMM approach is relatively computationally expensive and parameter dependent, and like any of these techniques will require adaptation and tuning for individual use cases. Fundamentally, by its design, MoSeq provides an alternate approach to the extraction of behaviours, defining putative behaviours directly from the state transitions modelled by the HMM rather than from the structure of the behavioural space.

### Discovering Behaviour for Comparative Neural Analysis

In this section, we will provide a detailed explanation step-by-step of how HUB-DT generates discovered behaviours from input data and enables comparative analysis with neural data, as well as our motivations for using the specific algorithms and techniques that make up the tool.

#### Extracting Pose: Annotating limb position from multi-camera recordings

The pipeline begins with data collection, to provide input for behavioural discovery. Single or multi video camera approaches are common across many of the tools in the field. Alternatives are depth cameras (or other depth measuring devices such as multiple infrared beams) or physical tracking where some sensor is placed on the animal. Each recording modality presents its own practical concerns in terms of experimental procedure and enclosure design. For our use cases, we utilize video cameras to allow simultaneous electrophysiology recording. This raw video is then annotated using the now field standard DeepLabCut (Mathis et al 2018): a supervised method for markerless tracking of lab animal features of interest from video (Figure 1: A-B).

**Figure 1:**
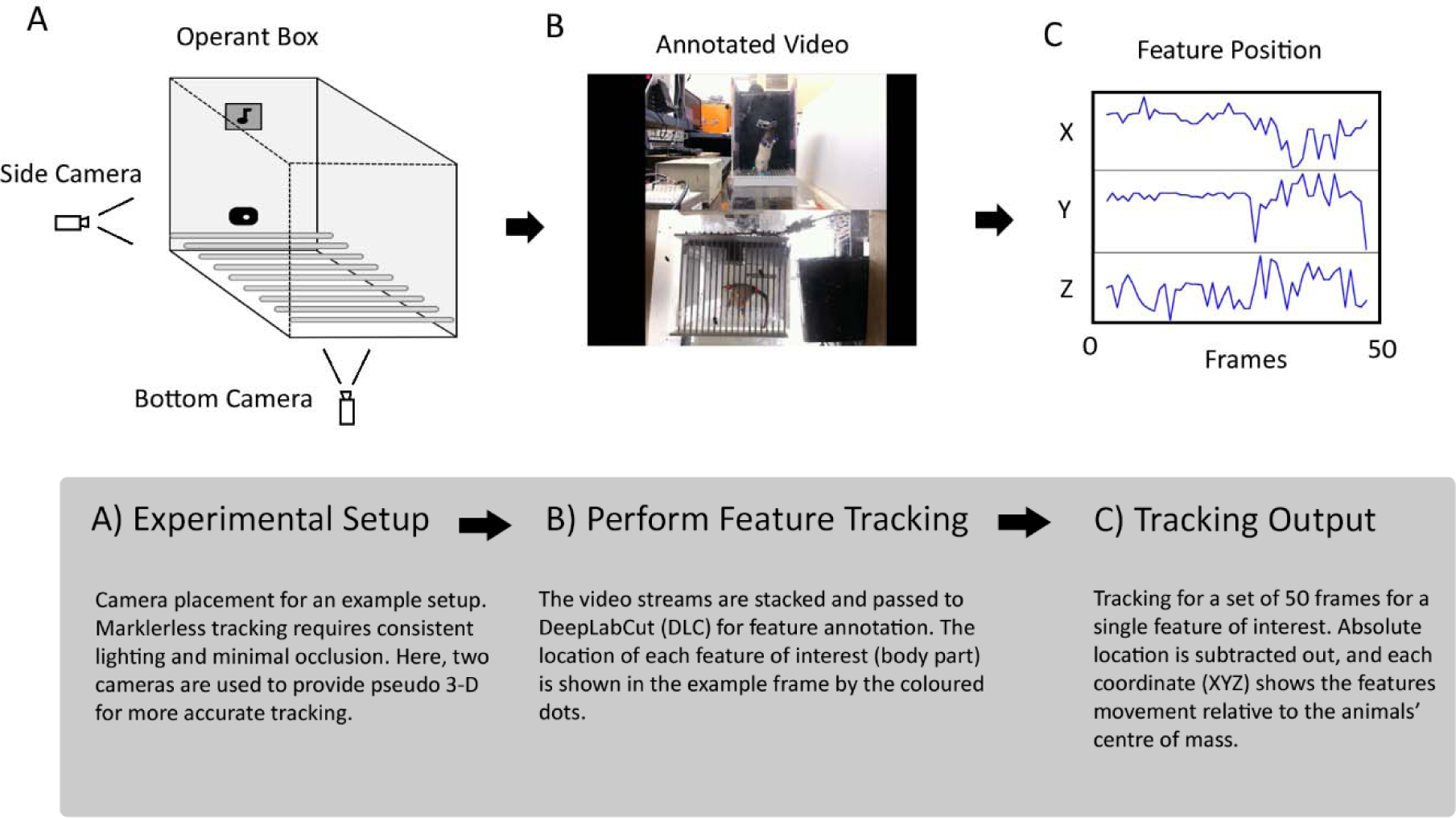
The HUB-DT Pipeline, A-C.

#### Behavioural Space: Projecting extracted pose into a spatiotemporal manifold

The output of DeepLabCut provides frame-by-frame positions (in 3-d) for our features of interest (Figure 1: C). As previously discussed, instantaneous body part locations (often collectively referred to as pose) does not in and of itself constitute behaviour. Rather, by our definition, behaviour is changes in pose that evolve over a range of timescales. Therefor, in order to capture behaviour both position and time components are necessary, specifically time components at multiple scales. Traditional fixed time period methods (like those used in many of existing tools in the field) have several key shortcomings. This is discussed at length by Berman et. al., in their Motion Mapper paper, but here we will cover them briefly. Notably, fixed time period inputs restrict temporal information to a specific scale. This discards multi-scale interactions across and within features that are present in many ‘typical’ behaviours. For example, rodent grooming behaviour is often indicated by paired high-frequency movements in the forelimbs, while the positions of the hindlimbs and other body parts like the snout and nose evolve more slowly. To completely describe these interactions, temporal information needs to be captured at multiple scales. This does not imply that fixed temporal scale methods are unable to categorize behaviours like grooming, but rather that the descriptions those tools generate of those behaviours may be incomplete. To capture motion at multiple temporal scales HUB-DT uses spectrographic projection via morlet wavelets to capture and discretize spatial and temporal features together in an expanded behaviour space, similar to Motion Mapper (Figure 2: D). Morlets, which are in essence complex gaussians, bias the space towards periodic movements, like those found in many expected behaviours such as locomotion, grooming, etc.

**Figure 2:**
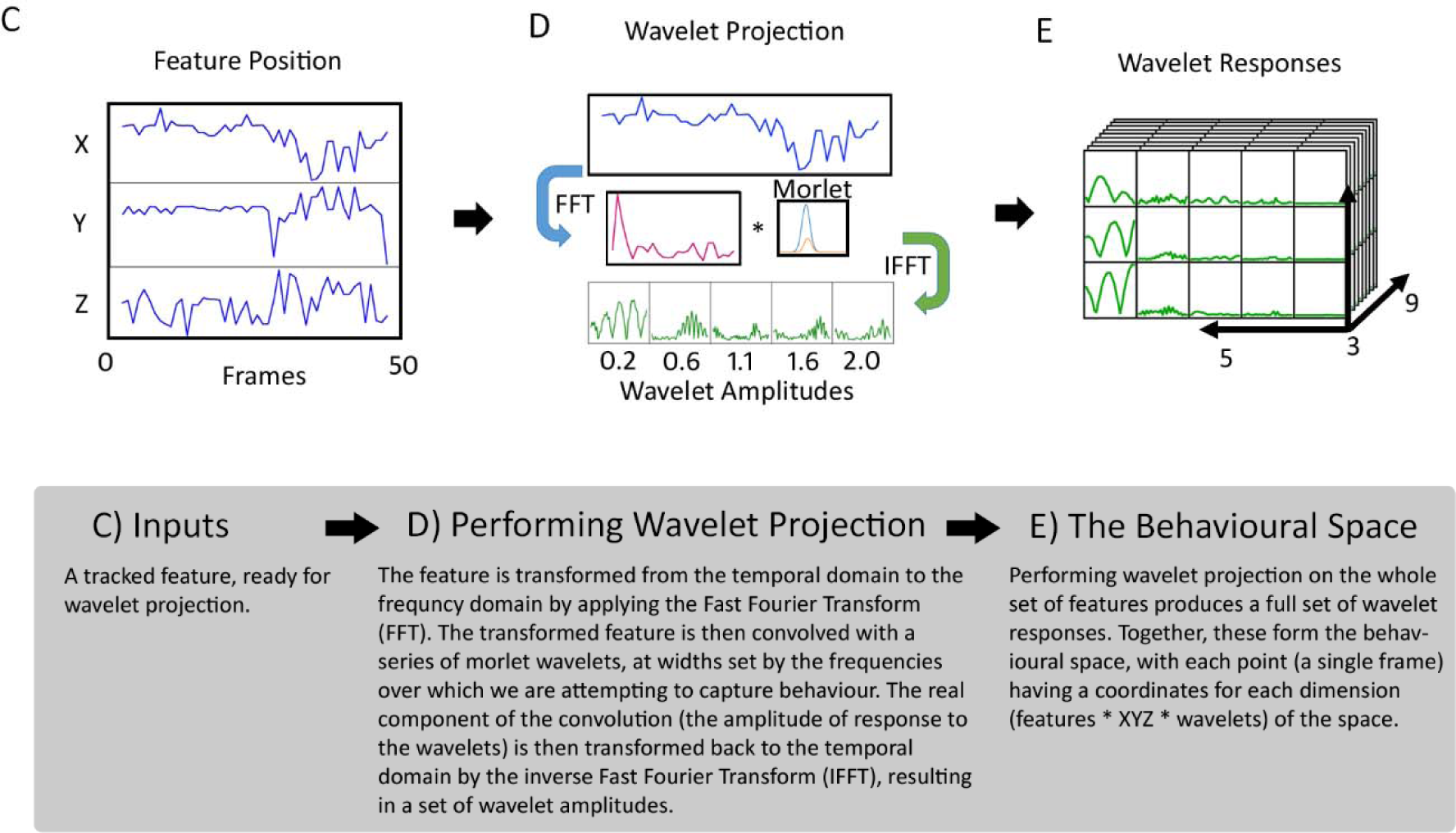
The HUB-DT Pipeline, C-E.

The end result of wavelet projection is a dataset where each point, associated with a single frame of video, is embedded in a high dimensional spatiotemporal manifold. The dimensions of this manifold are the response amplitudes of each of the features (and each of their 3 coordinates) to each of the wavelets used in the projection. The exact dimensions depend on the number of features and wavelets chosen, but for the pictured example the resulting manifold is 5 (wavelet frequencies) by 9 (features of interest) by 3 (tracked coordinates of each feature) (Figure 2: E). This space provides a complete description of the motion of tracked features, but its high number of dimensions renders it unsuitable for the application of many unsupervised clustering methods: this is due to both computational efficiency concerns, as well as to the ‘curse of dimensionality” which reduces algorithmic efficacy due to the burden of representing possibly uninformative dimensions during learning (Hastie et al, 2011). Complex learning algorithms like neural networks partially overcome this shortcoming by learning latent space representations of the data within the hidden layers, but in the interest of interpretability and computational resources we opted to perform dimensionality reduction prior to performing clustering.

#### Dimensionality Reduction and Density Maps

Dimensionality reduction is often carried out as a first pass in data pre-processing via Principal component analysis which yields a sequence of uncorrelated projections (linear approximations) of input data ordered by variance. A top subset of these projections can be used as a lower dimension approximation of the data preserving the bulk of the information. This reduced space is useful in identifying global attributes and trends in the data, but is not necessarily appropriate for cases where the information of interest is contained in local distributions of variable density, as is the case for our behavioural space data. There are a number of dimensionality reduction methods that are good candidates for preserving the information in the behavioural space, and here we cover a few that are implemented in our pipeline.

##### t-SNE

An alternative to simple component analysis is to use an embedding method like t-distributed stochastic neighbourhood estimation (t-SNE). t-SNE produces a lower dimensional (often as low as 2 or 3) embedding of the original data preserving proximity in the reduced space. This is achieved via modeling similarities in the input and output spaces as probability distributions (a series of gaussians in the input, and a t-distribution in the output). The distribution choice biases the embedding to preserve local structure (density, multiple scale relationships, manifold membership) in the data at the expense of global verisimilitude (as compared to other embedding methods that bias the resulting embedding in other ways). Base t-SNE is parameterized by a single parameter, perplexity (a continuous analog to k nearest neighbours) which defines the set of points over which the algorithm will attempt to preserve distances.

t-SNE’s suitability to producing low dimension embeddings of behavioural space is explored at length by Berman et al, in their Motion Mapper paper (Berman et al, 2014). Application on our rodent data was similarly successful, although there are some drawbacks with its use as part of a data processing pipeline. It is relatively computationally expensive (even on a modest data set) and is a transductive method, meaning it cannot be used to project new data into a calculated embedding. Much work has been done on improving t-SNE in recent years (van der Maaten et al., 2014, Linderman et al., 2017), and here we use the openTSNE package that utilises approximate neighbourhood calculations, interpolation, and multi-phase optimisation to greatly improve its performance, as well as providing a means for embedding new data. The structure of the embedding is visualised via a density map, showing the distribution of data in the reduced space.

##### UMAP

Uniform manifold approximation and projection (UMAP) is another embedding method which we can use to produce a lower dimensional (again for our purposes, as low as 2 or 3) embedding of the original data, as is demonstrated by Hsu et al, in their work on B-SoiD (Hsu et al, 2021). UMAPs’ approach shares some similarities with that taken by t-SNE, but follows a different path to optimize the match in structure between the data and its low dimensional representation. UMAP first constructs a ‘fuzzy topological representation’ of the input data, and this topology codifies the local structure of the data in its native high-dimensional form. Note, this summary statement conceals a swathe of algebraic topology methods which are used to justify and perform the construction of this representation (those interested in the process should refer to the UMAP publication: McInnes et al, 2018). Then UMAP attempts to optimize the fit (by cross-entropy) between a low-dimensional approximation of the topology with its high-dimensional original. Again, in practice UMAP utilizes a number of approximation techniques, and optimised initialisation steps to render the process computationally tractable. Similar to t-SNE, UMAP is parameterized by the number of neighbours used to calculate local structure. Adjusting this value changes the locality of the implied topology, essentially biasing the output to preserve structure at a particular scale. UMAP also allows control of the local density of the low-dimensional approximation via a minimum distance parameter. This has implications for downstream clustering, and allows for further fine-tuning of the output space. The UMAP package provides a scalable and relatively fast implementation, and again its output can be visualised via a density map of the data in the reduced space.

For practical purposes in constructing an analysis pipeline these two methods are often functionally equivalent, although care should be taken to account for each methods’ in-built biases, as mentioned above. In HUB-DT, the high-dimensional behavioural space data can be projected via either technique (or indeed other dimensionality reduction techniques) before being passed to further processing (Figure 3: F).

**Figure 3:**
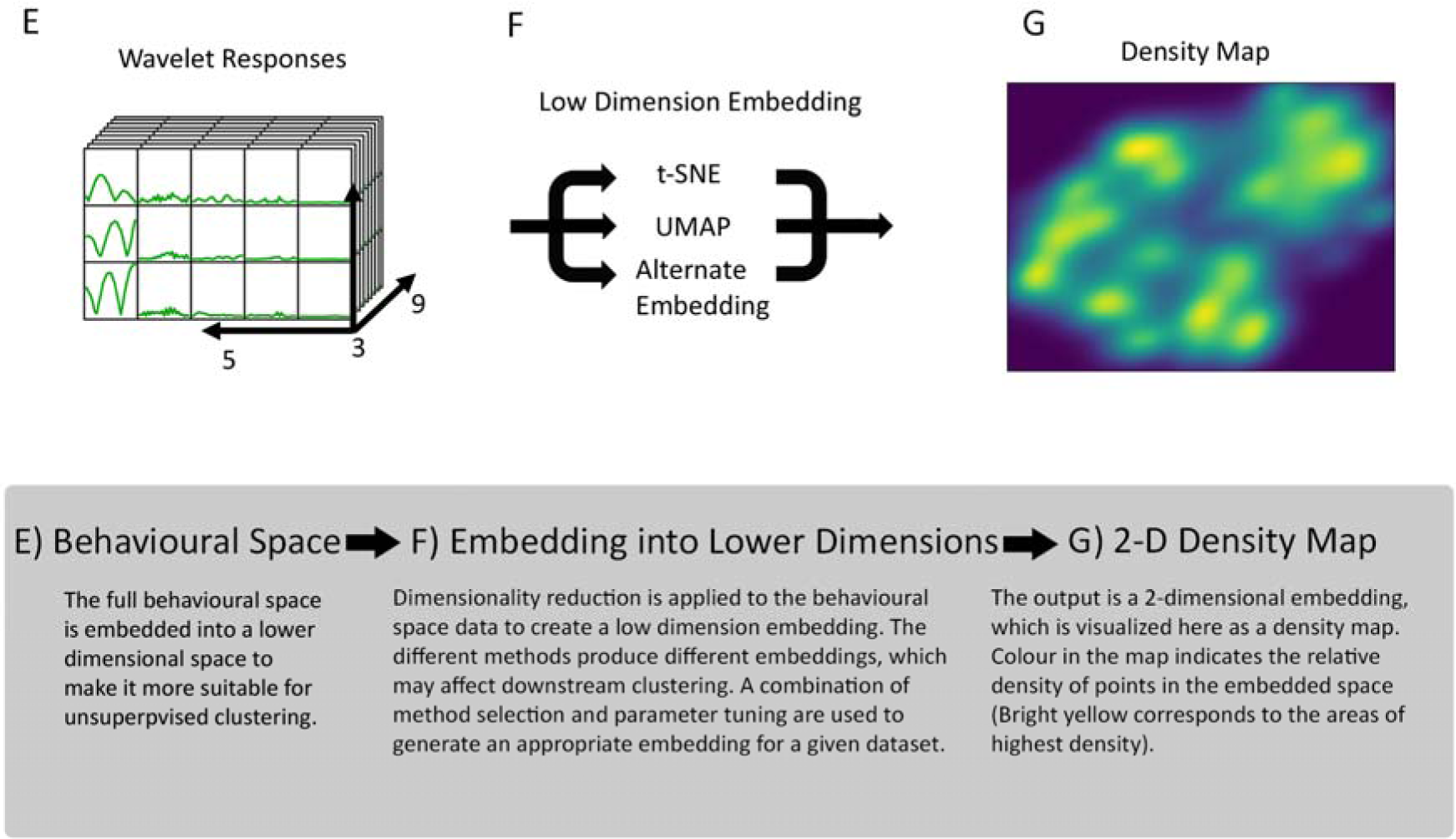
The HUB-DT Pipeline, E-G.

#### Hierarchical Unsupervised Clustering

The resulting density maps from performing dimensionality reduction on the behavioural space are useful for visualising the structure of behaviour across the whole dataset (Figure 3: G). Areas of high density show patterns of motion that occur frequently, and/or have low variance, ie motions that are highly stereotyped. Conversely, areas of low density correspond to motion patterns that occur infrequently, or have very high variance. We use this density landscape to extract structure from our data, segmenting the density map into clusters. This divides our data into a set of putative behaviours in an unsupervised way.

##### DBSCAN

Density-based spatial clustering with applications of noise (DBSCAN) is a flexible clustering technique that does not restrict clusters to be of any particular shape or distribution, rather clusters are separated from each other and from non-clusters via the concept of core samples (neighbourhoods of points in high density areas). Core samples form the core of clusters, and non-core samples that are within a given distance of a core sample form cluster fringes (distance here is a measure of relative density, defined as mutual reachability distance). Points further away are considered noise. Canonically, DBSCAN enforces clustering at a single density cut-off, the parameter epsilon, but by using a hierarchical approach DBSCAN can be extended to provide more complete and flexible cluster assignments.

##### HDBSCAN

A hierarchical extension of DBSCAN, as proposed by Campello et al (Campello et al, 2013). We use the HDBSCAN package (McInnes et al 2017) which modifies DBSCAN by constructing a minimum-spanning tree of reachability distances. This tree is converted into a hierarchy of connected points by iteratively merging points based on that reachability distance, producing a single linkage tree of the dataset. As opposed to a technique like DBSCAN, which slices this linkage at a flat value to produce a cluster labelling (any lone points above this value are labelled as noise), HDBSCAN condenses this tree with the use of a minimum cluster size threshold. Cluster splits that result in clusters containing fewer points than this threshold are treated instead as points ‘falling out’ of the parent cluster, ie labelled as noise. The result is a much-condensed tree, with clusters showing the persistence of structure in the data: clusters in the tree thin as points fall out and only split for sufficiently sized child clusters. This cluster persistence allows for the selection of clusters at variable density by working up the tree and selecting clusters of maximal stability to produce the final labelling. Of note, while HDBSCAN has the tendency to produce more clusters in the final labelling than most applications of DBSCAN, child-clusters are only selected if the sum of their stability/persistence is greater than that of their parent cluster, ensuring labelled clusters represent meaningful structure in the density landscape. Using HDBSCAN enables us to examine the hierarchy of clusters and gain some insight into the evolution of these clusters as we change the parameters of the algorithm.

**Figure 4:**
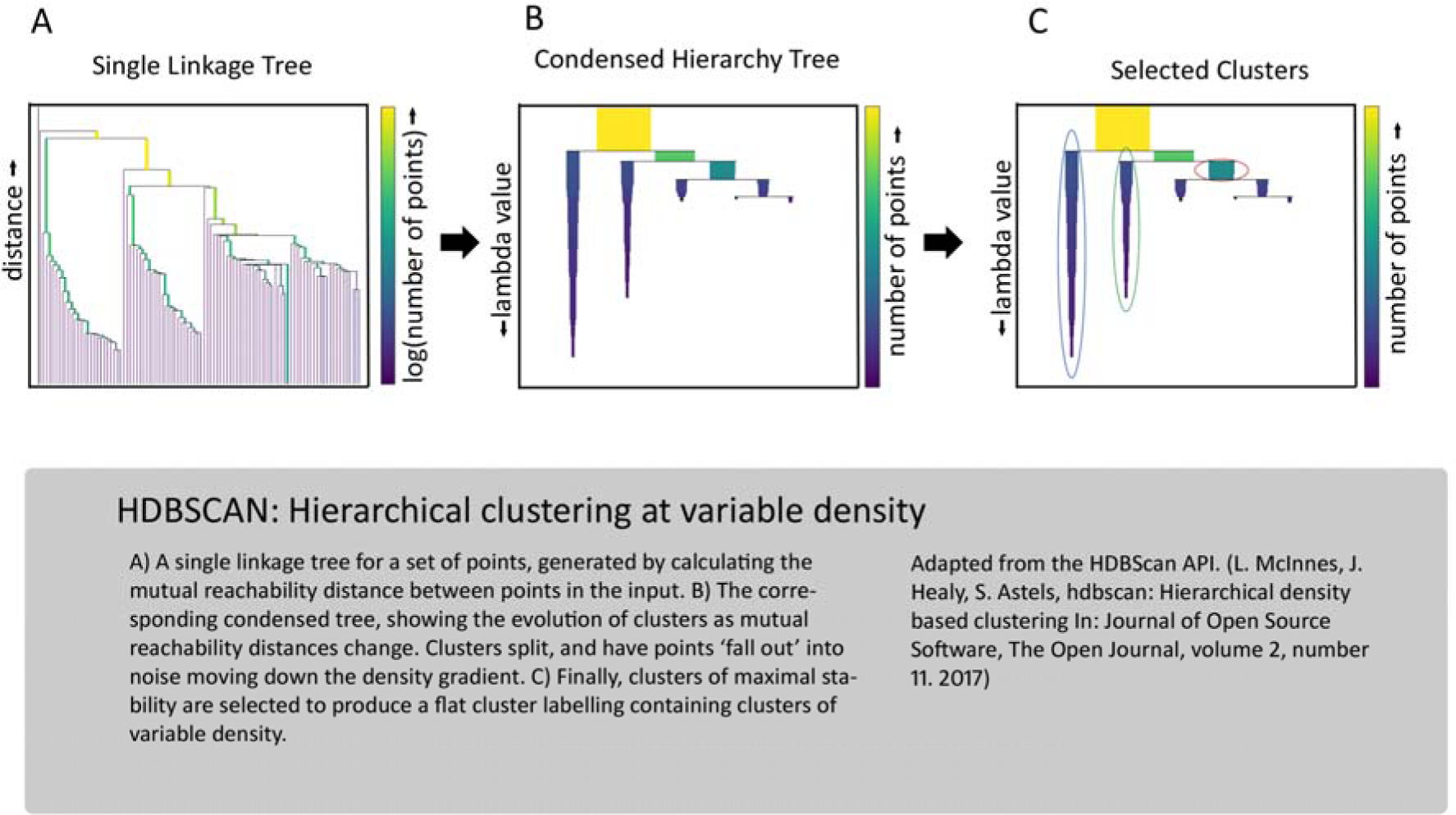
HDBSCAN.

**Figure 5:**
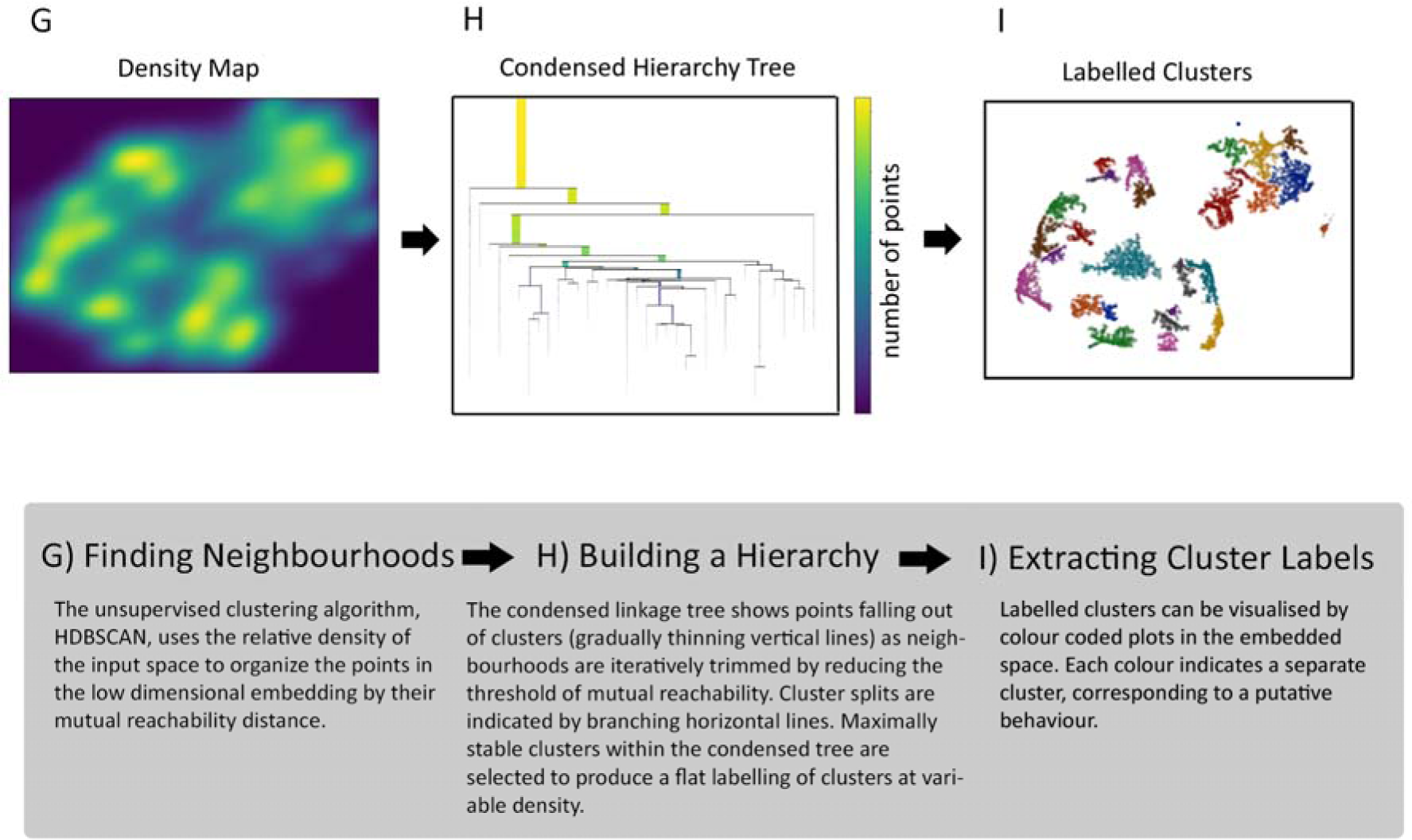
The HUB-DT Pipeline, G-I.

**Figure 6:**
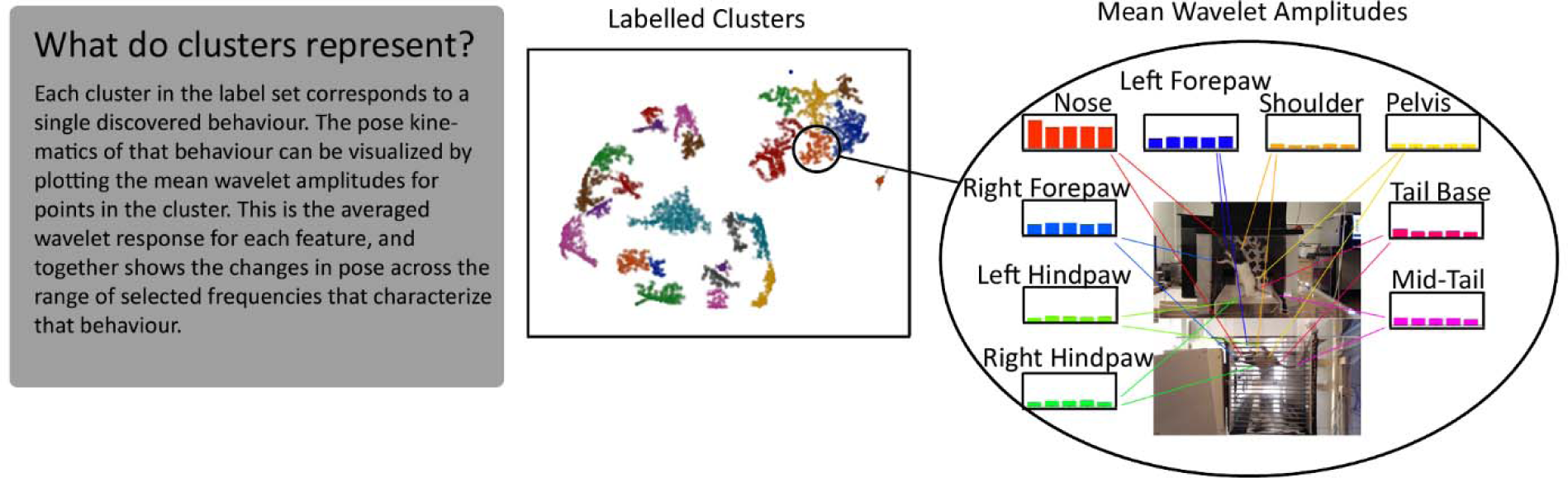
Clusters in Behavioural Space.

#### Connecting Unsupervised Behavioural Discovery to Manual Behavioural Labels

In an effort to make our unsupervised behavioural discovery more interpretable and relate it to manual behavioural annotation, we performed a series of supervised learning comparisons using the same behavioural space as defined above. Using our spatiotemporal projection as input, we trained a series of ensemble models to reproduce the manual labelling of specified behaviours such as locomotion, rearing, grooming, etc. To learn the human-annotated behavioral labels, RF’s were trained on a set of manually defined behavioural categories: inputs were samples from the full, high-dimensional behavioural space, and outputs were a set of binary vectors for each category. Each forest consisted of 100 trees, each built on a random subset of input dimensions (to a maximum of the square root of the total input dimensions) and on a bootstrapped sample of data. Split criteria in constructing each tree were based on maximising information gain amongst the given set of inputs, and outputs were calculated as an average of probabilities across the forest. The trees were optimised and evaluated using unseen training data from outside the sample (referred to as “out-of-bag” error). In this case, the RF’s trained on manual behavioural labels were evaluated not just for their performance (accuracy in predicting held out or novel data) but also for the subset of input dimensions, or features, from the full behavioural space that were most informative in categorising behaviours.

RFs: Random Forests (RFs) are a popular ensemble method for regression and classification problems. A large collection of de-correlated trees is constructed and trained, and their average output is used as a predictor. Trees are able to capture complex interactions between many correlated inputs, and benefit greatly from averaging to reduce instability and reduce variance. As such, random forests have a number of key advantages, notably low bias on high variance inputs, and are particularly useful in large dimension data sets with (relatively) a low number of samples like our behavioural space data (Hastie et al. 2011).

##### Feature Importance

The relative importance of a given feature in predicting can be evaluated within a random forest by looking at the expected proportion of samples they contribute to in each tree and then averaging those results, giving us a mean decrease in impurity measure, or ‘feature importance’ measure. These feature importances are easily calculable (via the mutual information measure used to define splits in the trees) but suffer from two key issues: they are based on training data statistics only, and are biased towards features with higher cardinality (more unique values). These can be mitigated by instead using a permutation-based importance measure, calculated on a held-out test set. Each feature is in turn permuted and the accuracy on this held-out set is compared between raw and permuted features. While permutation importance addresses the key flaws of impurity-based measures of importance, it unfortunately does not overcome underlying biases stemming from tree/forest construction: namely, a preference for strongly correlated features (Strobl et al., 2007). These biases can again be mitigated by building conditional permutation importance measures, but this is a rigorous and computationally intensive prospect.

##### Shapley Values and Universal Explainers

An alternative is to evaluate feature importance with a model agnostic method, by building a universal explainer; defined by Lundberg et al., 2017 as an interpretable approximation of the original, complex model. If properly constructed and defined, this explainer model can evaluate the contribution of features to a given output efficiently and agnostic of a given models particulars (Lundberg et al., 2017). This explainer can be used to calculate Shapley (SHAP) values as a measure of feature importance, again by a process of conditionally permuting the input and output space of the original model. Here we used the set of calculated SHAP values to extract the features of the full behavioural space that were important in our trained forests’ predictions of manually defined behavioural classes.

This set of importance features of the behavioural space defines a ‘human annotation’ manifold within the full behavioural space. By running our behavioural discovery process on this manifold, we can more directly compare the discovered behaviours to manually defined ones. In the restricted subset of dimensions, we made two key observations. First, that there are subsets of discovered behaviours that roughly align with manual behavioural labels, and second that manual behavioural labels typically align with multiple discovered behaviours, ie that the hierarchical clustering identifies structure within motion patterns associated with a manual behavioural label and breaks them down further into a set of related behaviours, or variations of the parent behaviour. By alignment, we are referring to similar structure in the density map, as defined by measures of comparative purity and cluster distance, which in turn indicates similar kinematics as measured by the average mean wavelet amplitudes of manual and discovered clusters.

Critically, when we expand the input to the behavioural discovery process to the full behavioural space, both of these observations remain true. HUB-DT is able to discover subsets of behaviours, amongst the full set of discovered behaviours, which are related to manual behavioural labels, but subdivides those manual categories. These similarities were assessed by the measures listed above, and additionally assessed qualitatively, by viewing video examples generated from the cluster labels. This subdivision separates out behavioral variations that are unlikely to be reliably recorded by a human observer, and in application have been found to be a more informative match for experimental conditions and neural data (this is discussed in more detail in the specific use cases).

**Figure 7:**
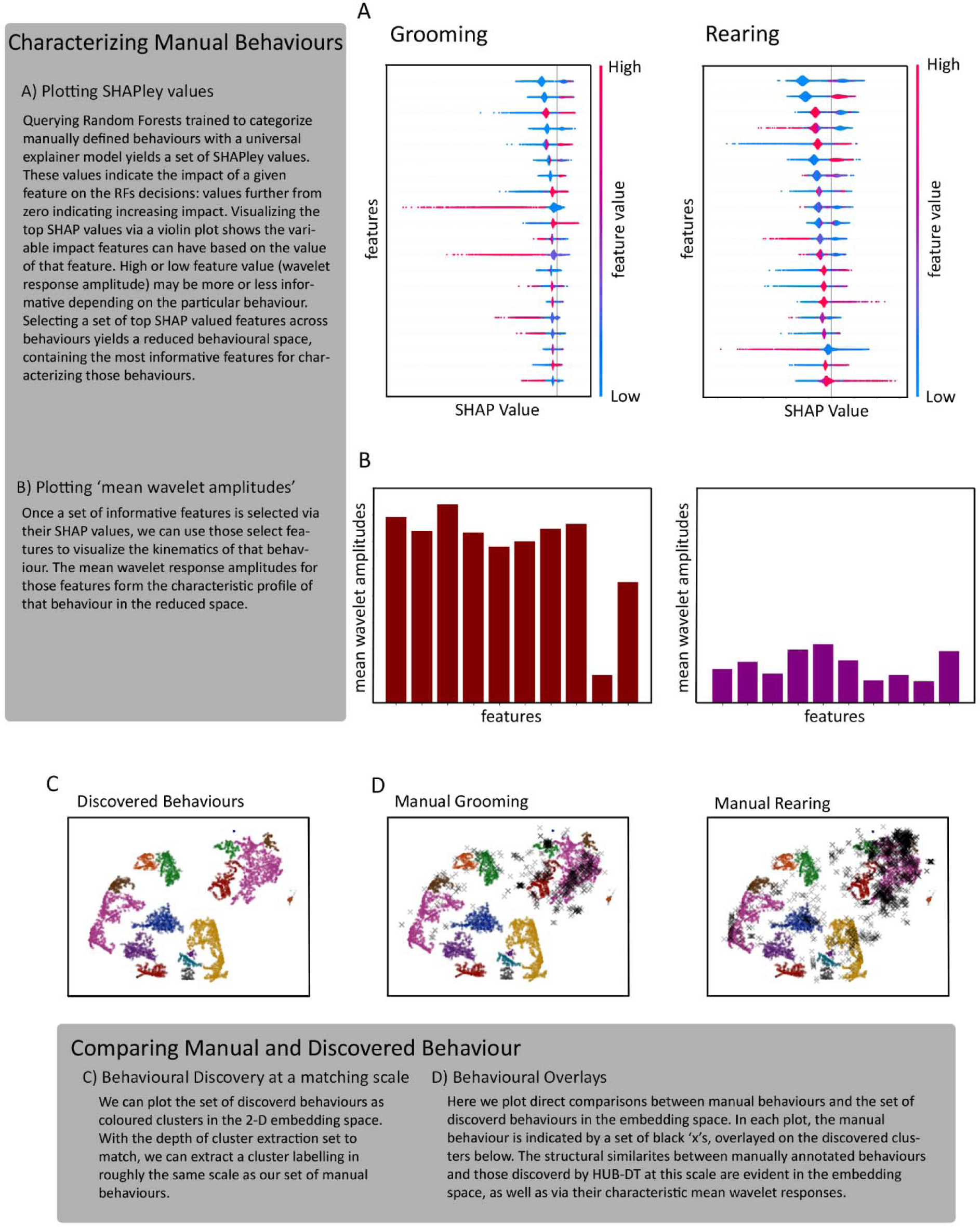
Characterizing Manual Behaviours.

**Figure 8:**
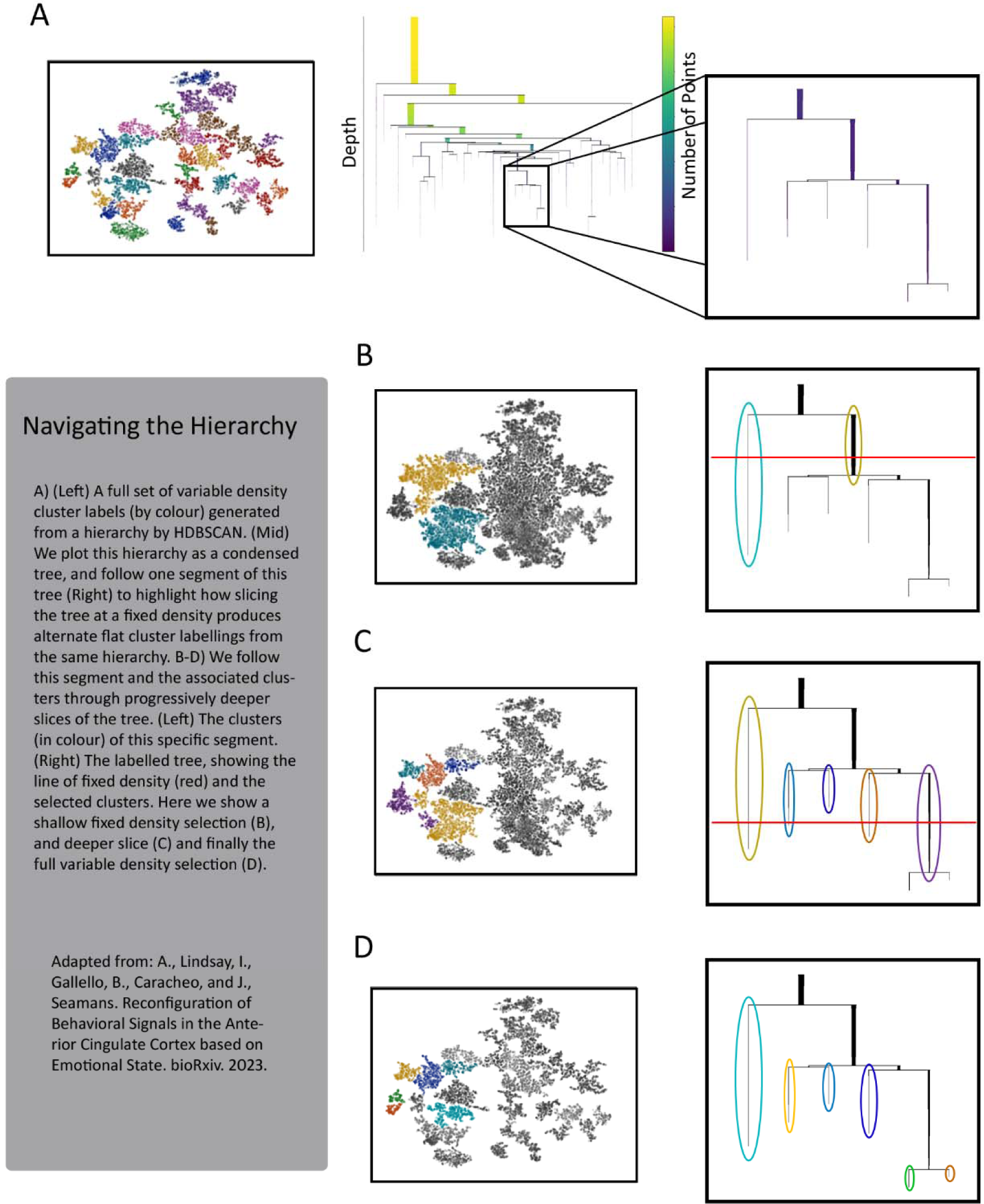
The Hierarchy of Behavioural Clusters.

#### Exploring the Hierarchy

As discussed, an advantage of a hierarchy-based clustering method is the ability to navigate that hierarchy and explore the structure and evolution of the behavioural clusters. By design, HDBSCAN assigns labels at variable density based on stable structures, but using the condensed tree we can also take fixed density slices to produce flat cluster labels at a variety of density thresholds. These slices help visualize how the specificity of behavioural labels evolves as we move deeper into the tree.

Sufficiently shallow slices provide only a rough separation of the behavioural space, and cluster structure can be as basic as simply separating locomotive behaviours from non-locomotive. Increasing depth in the tree is accompanied by increasingly specific behavioural clusters. This property means the cluster labelling, and therefor the selected behaviours, can be tuned to a desired granularity; for example, to match the scale and specificity of a set of manually selected behaviours. Or, can be left as a variable density labelling, to provide a completely unsupervised set of behaviours.

#### Evaluating discovered behaviour

Hierarchical clustering yields a set of defined clusters and a complete labelling of the set of frames into those clusters or un-clustered noise. The quality of the behavioural clusters identified are evaluated via a series of cluster metrics built in to HUB-DT. This includes direct cluster metrics: membership confidence, dwell time, and transition frequencies as well as behavioural measures such as mean wavelet amplitudes and variance within and across each cluster.

##### Adjusted Mutual Information and RAND Index

Mutual Information and the RAND index provide two statistical measures for comparing the fits between sets of labels. Mutual information is the measure of dependence between two variables, ie how informative one variable is of another. In comparing clusters, standard mutual information needs to be adjusted to account for differences in the size and number of clusters in the two sets, achieved by correcting the information measure for chance agreement. Similarly, the RAND index is a measure of the agreement between two sets of clusters. In essence it represents the frequency of occurrence of agreements over the sets of clusters, or equivalently the probability that a randomly chosen pair of points in one set of clusters will agree with the same pair in the other cluster set. The RAND index must also be corrected to account for differences in cluster size and number, by correcting by the expected similarity using a random model as comparison.

##### Split-Join Distance (Van Dongen Metric)

This distance is defined on the space of partitions (roughly equivalent to clustering) of a given set. It is the sum of the projection distance from one partition to another, and vise versa. Equivalently, it can be thought of as the number of steps required to rearrange one set of clusters into another, which each step being the relocation/relabelling of a single point. This measure is asymmetric, and so must be calculated for each direction. The van dongen metric provides a measure of the structural similarity of two sets of clusters, and as such may be more informative of good clustering fits on hierarchies, or other highly structured partitions compared to simpler measures of mutual information or precision/recall.

##### Fowlkes-Mallow Index (FMI)

the FMI is a score used to compare the quality of a match between two sets of labels. The index is defined as the geometric mean of pairwise recall and precision between the two sets. Values range from 0 to 1, with 1 being a perfect match between labels. In practice, it is often used to compare clustering generated labels to ‘ground truth’, but it is also useful for comparing hierarchical clusters with different structure.

##### Purity Measures

luster agreement can also be measured by a simple two-way cluster purity measure, calculated between two sets of labelled clusters. For example, if one set of cluster labels was for HUB-DT generated behaviours, and the other was for an experimental task factor. The purity measure would be the ratio between the proportion of a given behavioral cluster that was contained within a particular task factor cluster versus the proportion of the task factor cluster that was accounted for by the behavioral cluster. This purity measure is preferable to more complex statistical measures when the two sets of labels are expected to possess significantly different structure, or are incomplete, as in the given example.

The labelled clusters are further evaluated visually by inspecting frames and video of the putative behaviour. HUB-DT extracts the frames associated with each cluster and produces short videos of each behaviour. This aids in the interpretability of the discovered behaviours, as well as allowing for comparison between the statistical measures of cluster quality and the visual appearance of the behaviour.

**Figure 9:**
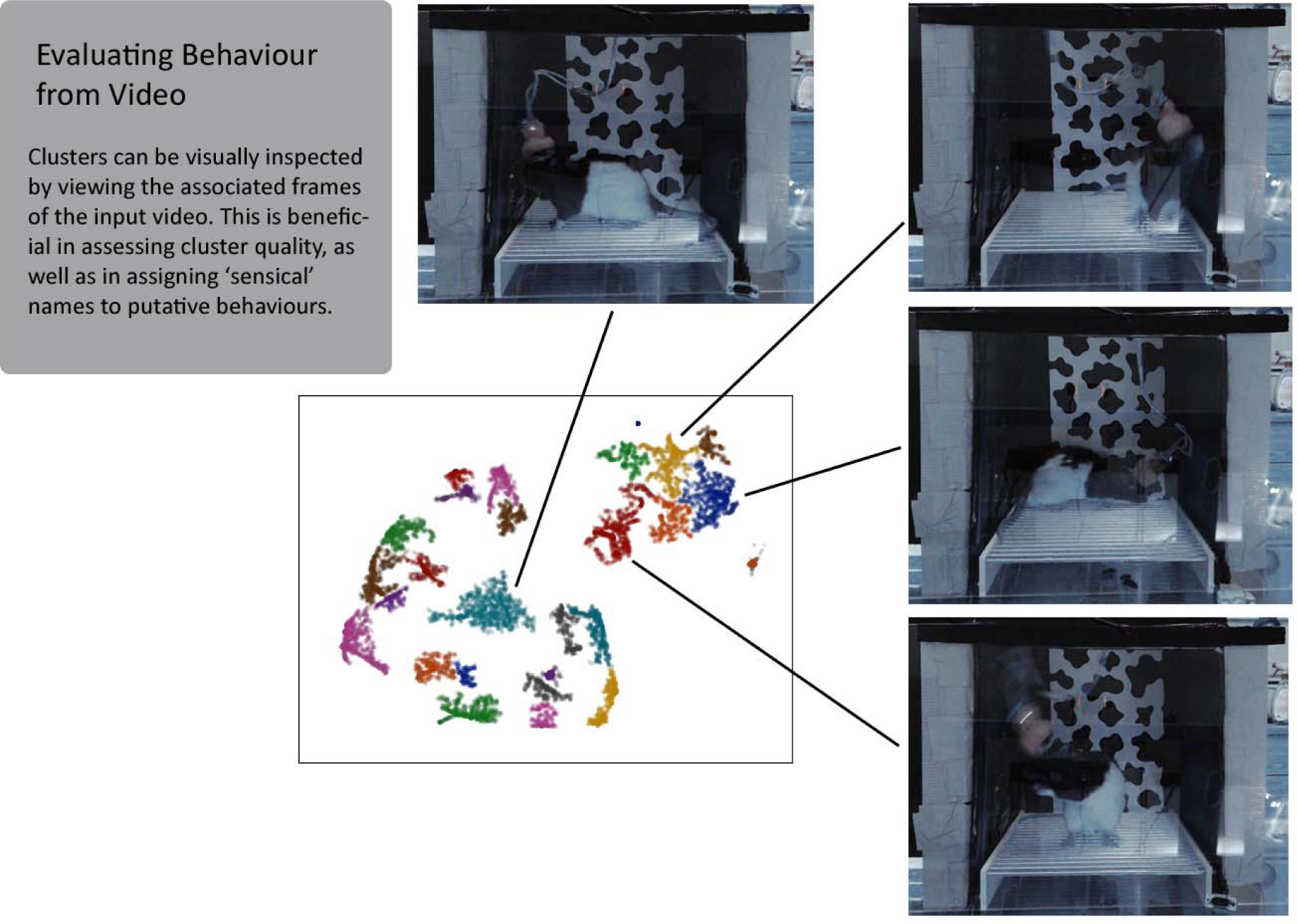
Visually Evaluating Behavioural Clusters.

This set of putative discovered behaviours can be propagated across additional datasets by the use of a trained supervised learning model. We use a Random Forest to learn the relationships between the full behavioural input space and cluster labels, and this trained model is then used to predict behavioural labels for new data, given the new data’s behavioural space as input.

### Comparative Analysis and Discovering Relevant Behaviour

We now have a set of putative discovered behaviours, and an accompanying labelled set of frames. It is worth, at this juncture, restating our operational definition of behaviour: sequences of pose over a range of timescales (this range is determined by the set of wavelets used in the spectrographic projection). These behaviours are not defined according to the animal’s goals, and may transition in timescales that are independent of the framerate at which the labels are set. While each frame is assigned a behavioural label, behaviours are not defined as transitions between poses in adjacent frames. Each instance of a given behaviour corresponds to a single point within that behaviour’s cluster that is defined across the whole range of timescales. This also means that each labelled frame contains information about changes in pose beyond the length of that single frame.

This labelled set of frames can now be used in comparative analysis with neural data and as a measure of the quality and neural relevance of discovered behaviour. In evaluating the efficacy of HUB-DT, we used paired neural data: electrophysiology data, recordings of multiple single neurons, binned at a matching framerate to the behavioural labels in a series of comparisons. In addition to utilizing the cluster metrics discussed above to evaluate neural clusters, unsupervised or defined by experimental factors, as compared to behavioural clusters, we also built statistical models to directly evaluate the relationship between neural data and discovered behaviour.

GLMs: The generalized linear models (GLM) used in HUB-DT take as input either neural data (usually time-binned recorded firing) as a Poisson sequence or behavioural space input with a selectable link function to a set of labels (such as HUB-DT generated behavioural labels, or imported task factors). GLMs can be single or multi-factor models, modelling the relationship between inputs and categorical factors. The strength of the relationship between a factor and input in the model is measured by the r^2^ value, the proportion of variance explained by a particular factor.

Comparative analysis can be performed with a whole range of statistical methods and machine learning models built into HUB-DT. These labels can also be exported for alternate analysis. The full scope of comparative analysis is covered in more detail in the discussed use cases, as well as in the referenced publications. As discussed above, comparative analysis is critical in evaluating appropriate behavioural clustering. The behavioural discovery process is iterative, using neural data to provide feedback for parameterization, adjusting the specifics of projection, dimensionality, and depth without the need for a priori definitions. Ultimately, this paired analysis yields a set of labelled behaviours with a measurable relevance to areas of interest within the brain.

### HUB-DT: Hierarchical Unsupervised Behavioural Discovery Tool

HUB-DT has been built into a python package for integrated behavioural discovery and neural analysis. The package is available on GitHub:

https://github.com/Loken85/HUB_DT/

The tool includes a GUI built using Streamlit (https://streamlit.io/). This GUI allows for interactive data exploration and analysis inside the browser, making it compatible for any operating system. Analysis is performed in real-time, and runtimes are reasonable even on low-end systems. A full run of behavioural discovery including computation should take no more than ∼10 minutes, even for long (hours) experiments. More computationally intense operations, like training supervised learning models, can take longer, but these models can be saved and exported/imported for re-use.

Here we have included a quick user guide for the HUD-DT application. This guide covers the basic layout of the GUI, and provides an example workflow for running behavioural discovery and some basic combined analysis using HUB-DT.

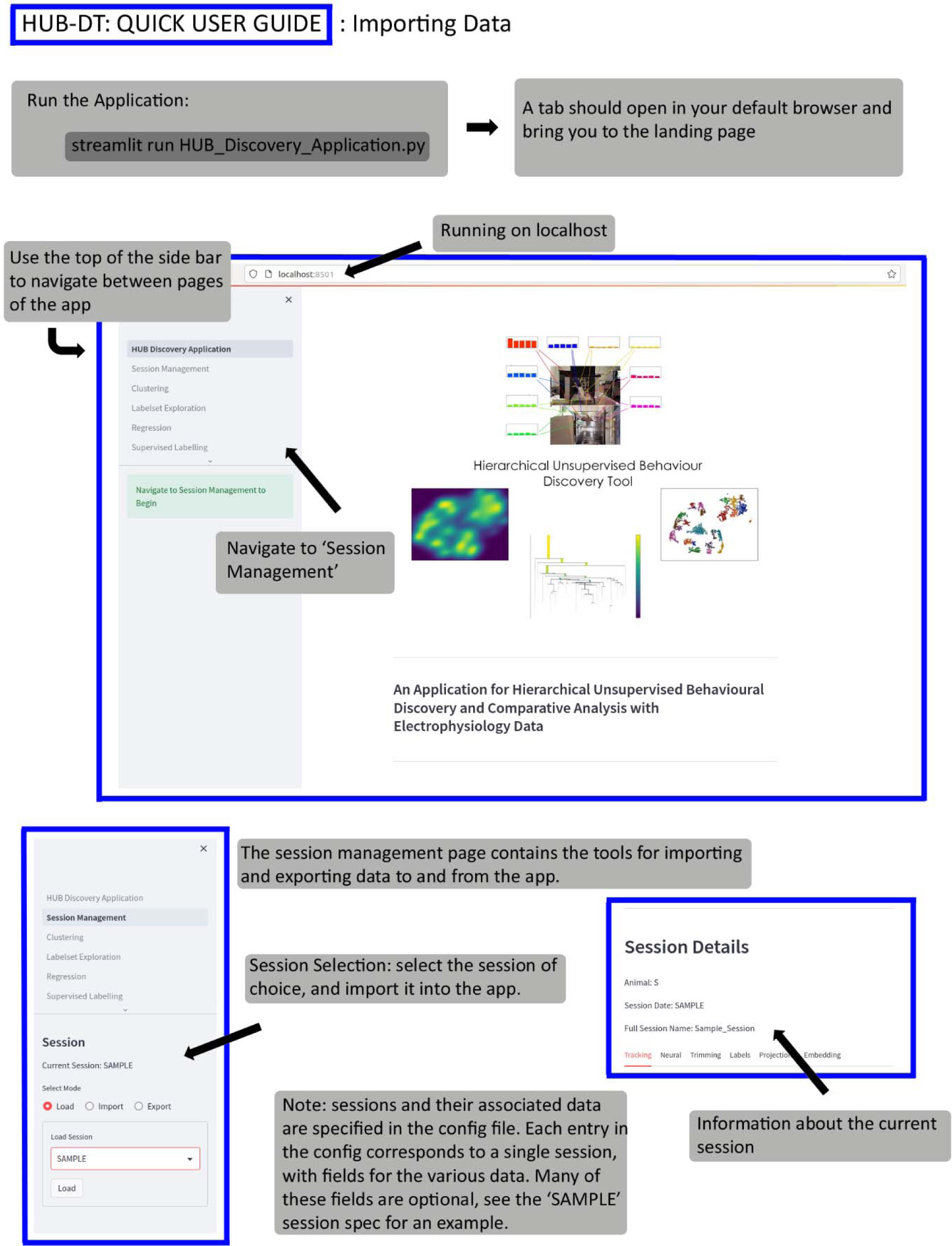
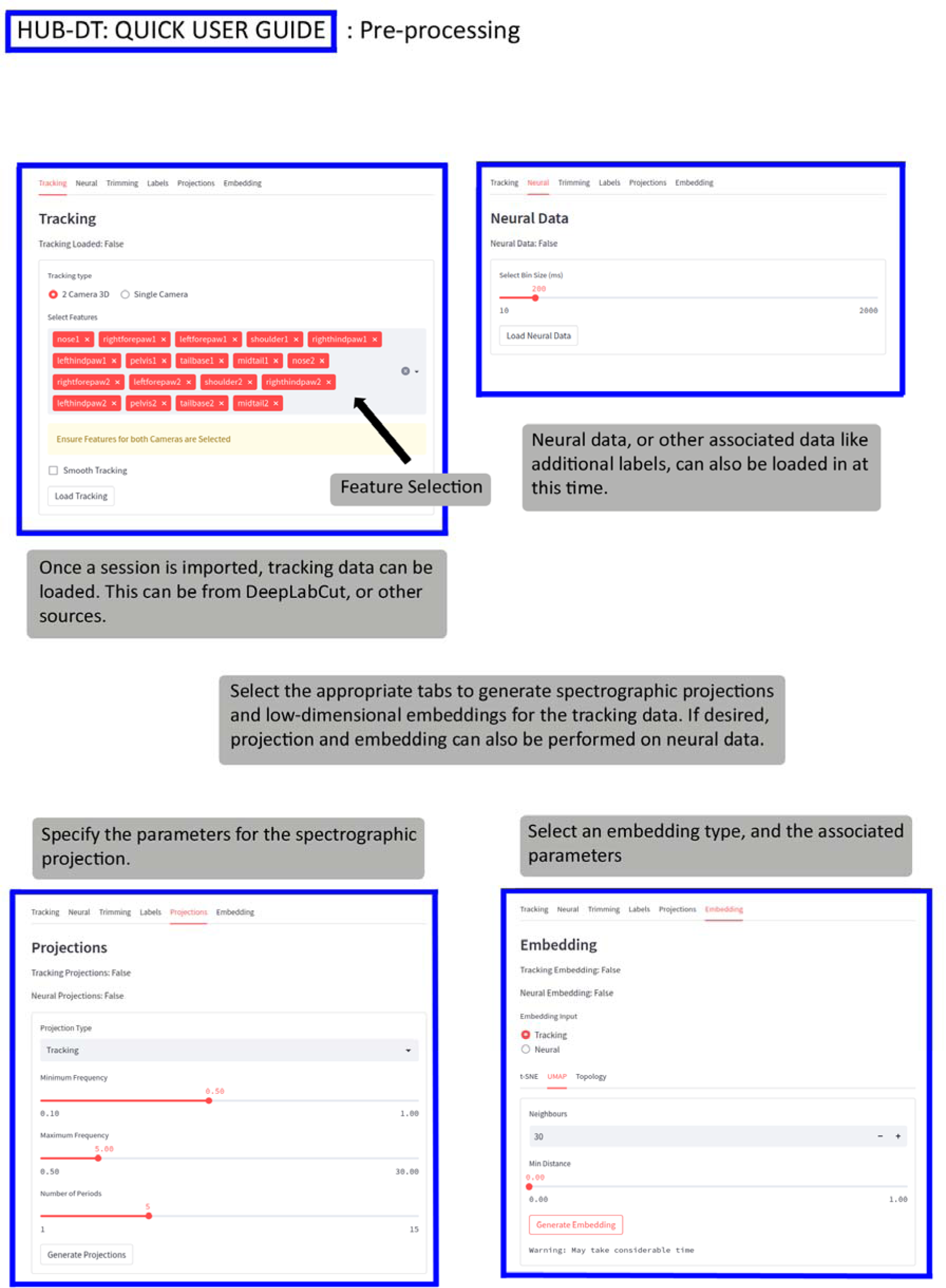
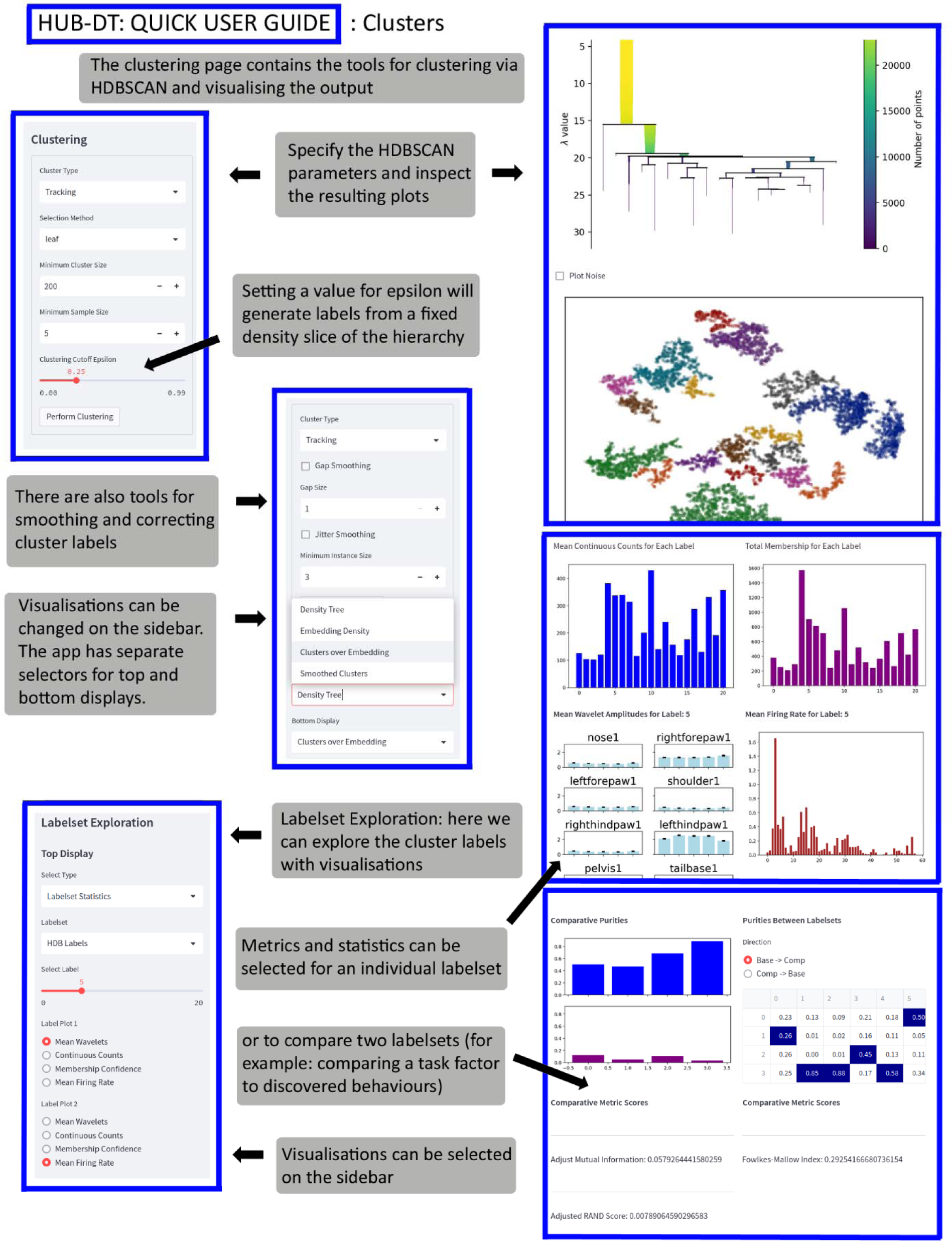
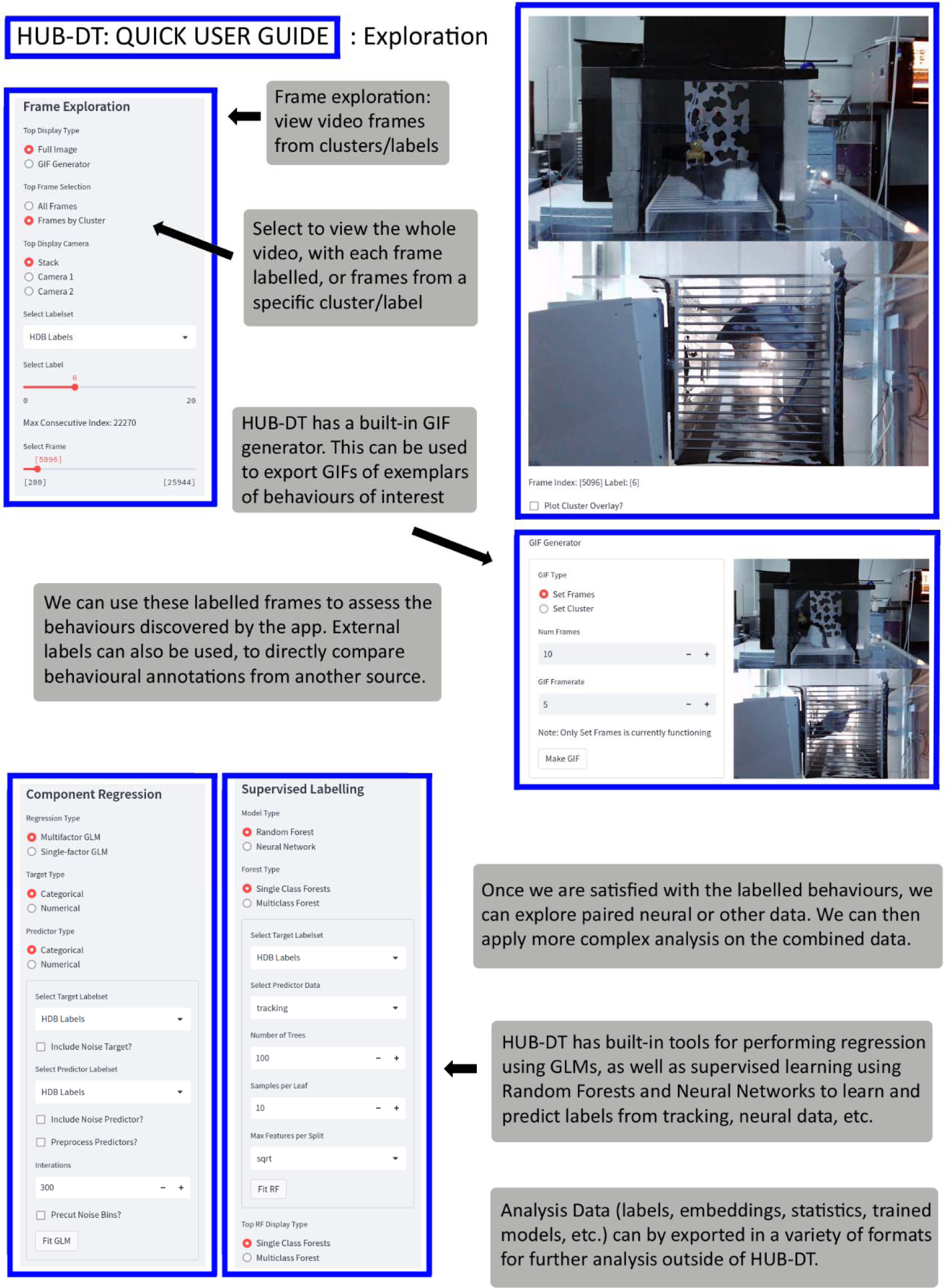

### HUB-DT Use Cases

In the following section we have compiled some use cases for HUB-DT from work in our own lab. These cases highlight the utility of the analysis pipeline across a variety of applications in our work, and represent a body of both published and in-progress studies. We will attempt to provide some context for the application of HUB-DT in each case, and briefly cover some insights that our analysis revealed.

### Investigating the characteristics of neural activity related to behaviors versus other task events

Our work is focused on the function of the frontal cortex, and in particular the anterior cingulate cortex (ACC). ACC neurons can flexibly multiplex a staggering array of information, which makes it incredibly challenging to understand what is driving any given neural response.

In an effort to understand how the neural representation of behaviours in the ACC was modified or modulated by changing emotional state, we applied HUB-DT to identify and categorize spontaneous behaviours during a task designed to elicit distinct ‘emotional contexts,’ and performed a series of neural activity analysis to identify and separate neural signals associated with behaviour and those emotional contexts (Lindsay et al 2023). The data were collected as rats performed our ‘3-valence’ task (Caracheo et al 2018), where tones were paired with outcomes having a different emotional valence. The task is structured into 3 blocks with each block being dedicated to trials to outcomes of a specific valence (food, foot shock, and neutral or no outcome). Each block is treated as a unique ‘emotional context’. We have observed that each of these emotional contexts is associated with a unique ensemble activity state pattern in the ACC that persists throughout the entirety of the block (Caracheo et al 2018; Lindsay et al 2023).

**Figure 10:**
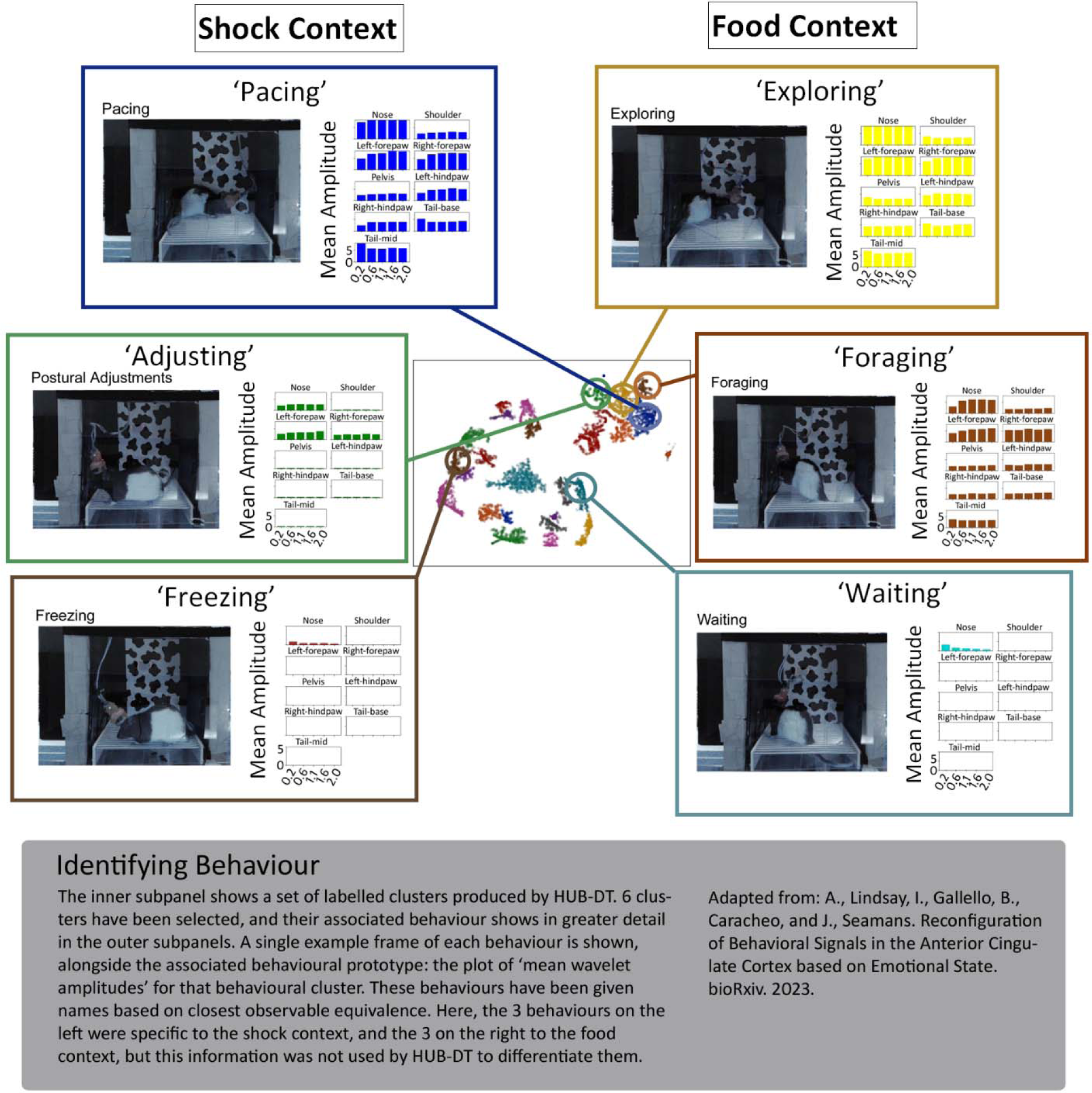
Identifying Emotional Context Specific Behaviours.

A possible issue with assigning emotional context dependent states is that as ACC neurons also encode actions and behaviors, the unique patterns we observed could have arisen not because the ensembles were tracking emotional states per se but because they were tracking the unique behaviors that are emitted whenever rats enter one of these emotional states.

In Lindsay et al (2023) HUB-DT was utilized to discover a large set of behaviours present during the task, including both some behaviours that were unique to specific emotional contexts, as well as behaviours that occurred across multiple contexts, and to separate neural signals related to behaviour versus the emotional context in which they are embedded. Many of these discovered behaviours were easily identifiable to human observers, and included expected contextually appropriate behaviours (such as freezing in the shock block and foraging behaviour in the food block) even though the behavioural discovery process was agnostic to the contexts and other task specifics (Figure 10). By applying paired analysis of this discovered behaviour with our electrophysiology data, we were able to identify the proportions of recorded neurons that were responsive to behaviour, and context, and highlight the distribution of multi-selectivity within the population. We demonstrated that roughly 10 times as many neurons responded to behaviours in a contextually dependent versus fixed manner, and that those behavioural representations appear to be embedded within the overall context. These context representations form distinct clusters in neural space, but despite the dramatic shifts between contexts the representations of individual behaviours remain identifiable.

**Figure 11:**
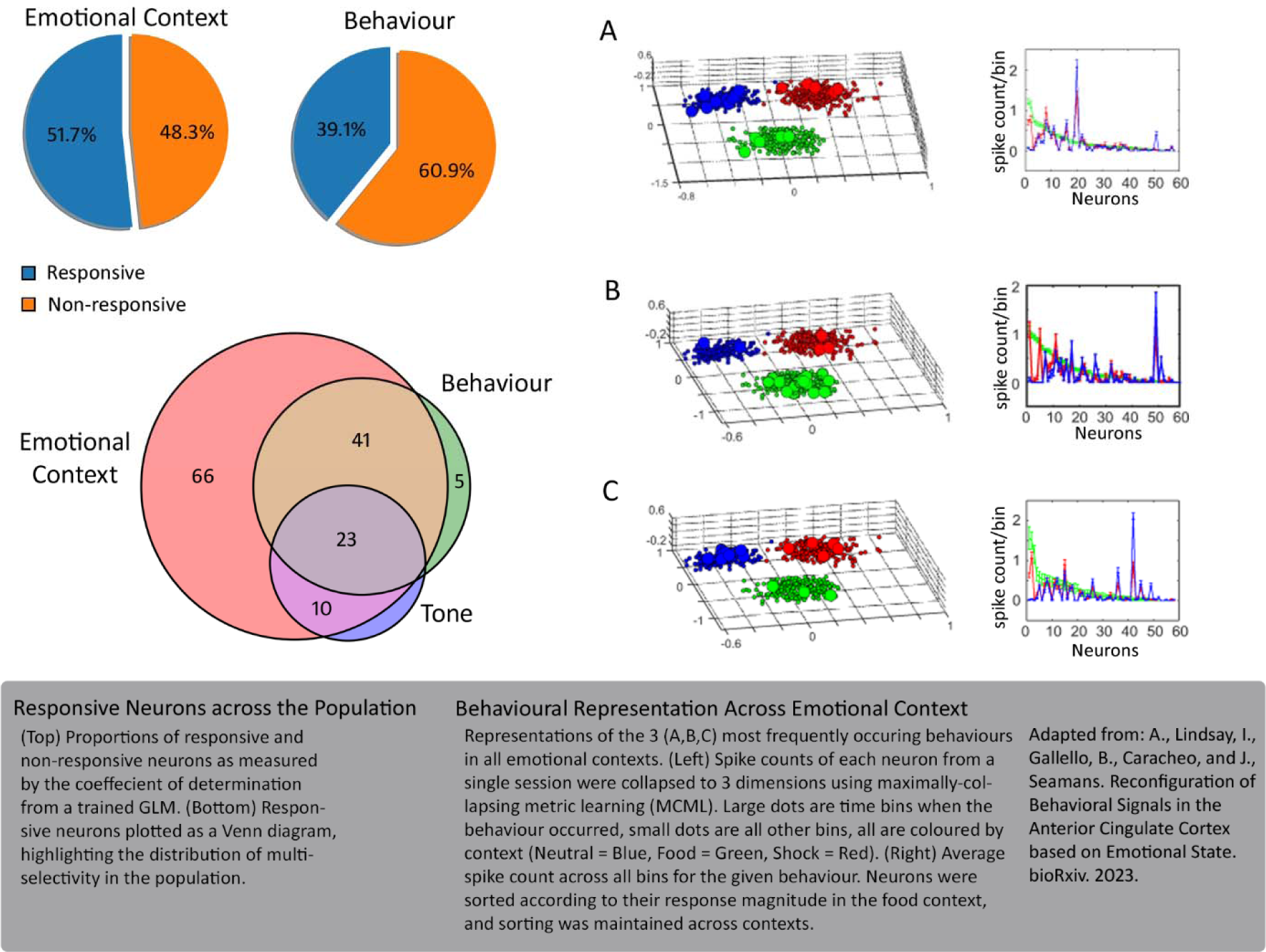
Neural Representation of Behaviour Across Emotional Contexts.

### Discovering the behaviours of addiction and withdrawal

Identifying behaviors evoked by a specific drug of abuse or stage of addiction have traditionally been based on the impressions of human observers. For instance, the behaviors that are typically associated with opioid exposure include: frozen postures, sniffing-up, grooming, rearing, confined gnawing, Straub tail and licking whereas the behaviors associate with opioid withdrawal include: jumping, rearing, paw flutters, wet dog shakes, abdominal constrictions, hunched posture, and teeth chattering (Gellert and Holtzman 1978; Maldonado et al. 1992; Maldonado and Koob 1993; Belozertseva et al 2016). There are some clear issues with this approach. First, while some of these behaviors, like Straub tail (prolonged tonic dorsal extension of the tail) are morphine-specific, many are not. Second, the presence or absence of any of these behaviors is at the discretion of the rater and their skill level. Third, even if a rater is highly skilled, the process is extremely time-consuming as the rater must advance through the video and score frame(s) manually. Automated behavioural detection and discovery alleviates many of these issues, and enables the labelling of potentially undefined behaviours that human observers would likely miss.

Here we applied HUB-DT to behavioural analysis in a morphine use and withdrawal study, with the goal of being able to reliably detect behaviours specific to drug states. For this study, rats received morphine on a 5 days on, 2 days off injection schedule. Each day they were placed in an open field box for 30 minutes and were recorded with 2 cameras, mounted below and to the side of the enclosure. DeepLabCut was applied to provide pseudo-3-D tracking.

**Figure 12:**
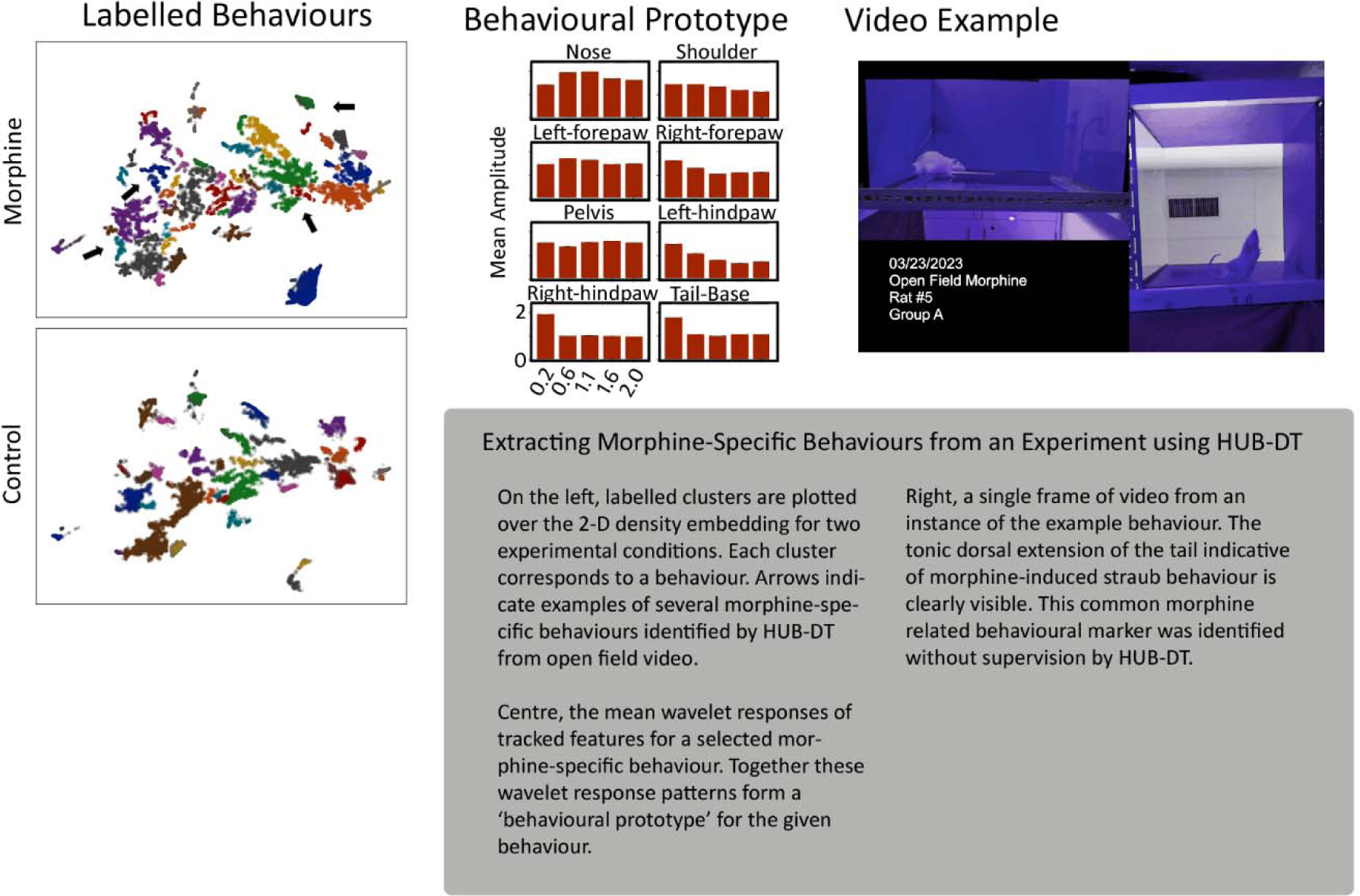
Extracting Morphine-Specific Behaviours.

HUB-DT was able to reliably identify several morphine-specific behaviours. These drug-state dependent behaviours were extracted purely from the spatiotemporal projection and hierarchical clustering utilized by HUB-DT, without the need for a priori definitions or supervision. Examination of these discovered behaviours, by inspection of the behavioural prototypes, as well as labelled video generated by the tool showed that they included consistently observed and commonly used markers of morphine related behaviour in rodents, such as morphine-induced Straub tail (Belozertseva et al 2016).

## Acknowledgements

This work was supported by funding from CIHR (MOP-137045 and PJT-159796) and NSERC (05979)

